# Inositol hexakisphosphate kinase 1 is essential for cell junction integrity in the mouse seminiferous epithelium

**DOI:** 10.1101/2023.04.26.538408

**Authors:** Sameer Ahmed Bhat, Aushaq Bashir Malla, Vineesha Oddi, Jayraj Sen, Rashna Bhandari

**Affiliations:** Laboratory of Cell Signalling, Centre for DNA Fingerprinting and Diagnostics (CDFD), Inner Ring Road, Uppal, Hyderabad 500039, India; Graduate studies, Manipal Academy of Higher Education, Manipal 576104, India

**Author notes:** Current address: Department of Biochemistry and Molecular Genetics, College of Medicine, University of Illinois at Chicago (UIC), IL 60607, USA. Current address: Department of Genetics, Yale School of Medicine, New Haven, CT 06510 USA. Current address: Syngene International Limited, Biocon Park, Bengaluru, Karnataka 560099, India (Vineesha Oddi). These authors contributed equally to this work (Sameer Ahmed Bhat, Aushaq Bashir Malla). Correspondence to Rashna Bhandari.

**Keywords:** IP6K, Blood-testis barrier, Cell junctions, Spermatogenesis, Actin cytoskeleton **Abbreviations:** IP6K1, inositol hexakisphosphate kinase 1, BTB, blood-testis barrier, ES, ectoplasmic specialization, dpp, days postpartum.

## Abstract

Inositol hexakisphosphate kinases (IP6Ks) are enzymes that catalyse the synthesis of the inositol pyrophosphate 5-IP7 which is involved in the regulation of many physiological processes in mammals. The IP6K paralog IP6K1 is expressed at high levels in the mammalian testis, and its deletion leads to sterility in male mice. Here, we show that the loss of IP6K1 in mice causes a delay in the first wave of spermatogenesis. Testes from juvenile *Ip6k1* knockout mice show downregulation of transcripts that are involved in cell adhesion and formation of the testis-specific inter-Sertoli cell impermeable junction complex known as the blood-testis barrier (BTB). We demonstrate that loss of IP6K1 in the mouse testis causes BTB disruption associated with transcriptional misregulation of the tight junction protein claudin 3, and subcellular mislocalization of the gap junction protein connexin 43. In addition to BTB disruption, we also observe loss of germ cell adhesion in the seminiferous epithelium of *Ip6k1* knockout mice, ultimately resulting in premature sloughing of round spermatids into the epididymis. Mechanistically, we show that loss of IP6K1 in the testis enhances cofilin activity due to increased AKT/ERK and integrin signalling, resulting in destabilization of the actin-based cytoskeleton in Sertoli cells and germ cell loss.

## Introduction

Inositol hexakisphosphate kinases (IP6Ks) are enzymes that catalyse the synthesis of the inositol pyrophosphate 5-diphosphoinositol pentakisphosphate 5-IP7 from inositol hexakisphosphate (IP6) [1, 2]. Mammals possess three isoforms of IP6Ks – IP6K1, IP6K2 and IP6K3, that vary in their tissue and subcellular distribution [3, 4]. IP6K1 is involved in many physiological processes including spermatogenesis, insulin secretion, vesicular trafficking, blood clotting, and cell migration [4-9]. Knockout of *Ip6k1* in mice causes male sterility [5]. One reason for this is the absence of ribonucleoprotein granules called chromatoid bodies in *Ip6k1* knockout round spermatids, leading to premature translation of the sperm-specific nuclear proteins TNP2 and PRM2 [10]. This results in faulty sperm differentiation and caspase-dependent apoptosis of elongated spermatids. Another reason for infertility in *Ip6k1^-/-^* male mice is abnormalities in the cell junctions that connect elongating spermatids with Sertoli cells in the apical region of the seminiferous epithelium [11].

The seminiferous epithelium in the testis is composed of Sertoli cells connected with each other, and with germ cells at different stages of their development, through multiple cell-cell junctions (Supplemental Figure 1A) [12]. These testicular junctions can be broadly divided into three types - tight junctions, gap junctions, and anchoring junctions. Anchoring junctions are of two types: (a) actin filament-based junctions, which include adherens junctions and focal adhesions, and (b) intermediate filament-based desmosome-like junctions [13]. In contrast to somatic tissues where cells at a particular location are connected by only one type of junctional complex, testicular junctions are heterogeneous - two or more types of junctions coexist in close proximity within the seminiferous epithelium [14]. The best example of testicular junctional heterogeneity is provided by the blood-testis-barrier (BTB), which is a junctional complex between two adjacent Sertoli cells (Supplemental Figure 1B). The BTB is composed of tight junctions, adherens junctions, gap junctions, and desmosome-like junctions, all of which are present together in the epithelium [15]. The BTB acts as an impermeable barrier, dividing the seminiferous epithelium into basal and adluminal compartments (Supplemental Figure 1A). This excludes post-meiotic germ cells present within the latter compartment from systemic circulation, preventing an autoimmune response towards spermatid antigens [12].

The ectoplasmic specialization (ES) is an atypical actin-based adherens junction present in the testis [16]. Two types of ES are present in the seminiferous epithelium - basal ES, and apical ES. The basal ES is present at the BTB between two adjacent Sertoli cells, whereas the apical ES is restricted to the Sertoli cell-spermatid interface towards the lumen of the tubule (Supplemental Figure 1A, B and C) [16, 17]. Disruption of the apical ES leads to spermatid polarization defects and immature release of spermatids from the epithelium [13].

In this study, we have examined the defects in different types of cell junctions within the seminiferous epithelium of *Ip6k1* knockout mice. Gene expression analysis show a downregulation of cell adhesion pathways in testes from juvenile *Ip6k1^-/-^* mice. Tissue immunofluorescence and immunoblotting assays reveal that the loss of IP6K1 in mouse testis causes gross abnormalities in cell adhesion within the seminiferous epithelium. We observe that IP6K1 deletion is associated with compromised BTB integrity and aberrant Sertoli cell-Sertoli cell and Sertoli cell-germ cell junctions. At the molecular level, we show that the absence of IP6K1 results in dysregulation and mislocalization of junctional proteins, as well as abnormal integrin and growth factor receptor signalling. These molecular and structural anomalies may be the underlying basis for seminiferous epithelium disruption in *Ip6k1^-/-^* testes, and contribute to the failure of spermatogenesis in these mice.

## Results

### The first wave of spermatogenesis is delayed in *Ip6k1^-/-^* mice

During the first wave of spermatogenesis in juvenile mice, developing germ cells advance synchronously through the seminiferous epithelium with the sequential appearance of spermatogonia, spermatocytes, round spermatids, elongating and elongated spermatids [18] (Supplemental Figure 1D). To identify the phases of spermatogenesis that are affected by the loss of IP6K1, we histologically examined juvenile testes from 14, 18, 24, 28 and 34 days postpartum (dpp) mice. At 14dpp, the seminiferous tubules were composed of Sertoli cells and spermatogonia, along with early pachytene spermatocytes. However, the tubules containing early pachytene spermatocytes were fewer in *Ip6k1^-/-^* compared with *Ip6k1^+/+^* testes, suggestive of a lag in meiotic initiation in the absence of IP6K1 (Figure 1, A and B). At 18dpp, when meiosis reached the late pachytene and diplotene stages in *Ip6k1^+/+^* testes, there was a significant reduction in tubules containing late pachytene and diplotene spermatocytes in the testes of *Ip6k1^-/-^* mice of the same age (Figure 1, C-E). At 24dpp, round spermatids were observed in the testes of *Ip6k1^+/+^* mice, but *Ip6k1^-/-^* testes showed fewer round spermatids (Figure 1, F and G). At 28dpp, meiosis was complete in both *Ip6k1^+/+^* and *Ip6k1^-/-^*testes, there being no significant difference in the fraction of tubules containing round spermatids (Figure 1, H and I). Early elongating spermatids marking the initiation of spermiogenesis were present in 28dpp *Ip6k1^+/+^* testes, whereas these germ cells were nearly absent in *Ip6k1^-/-^* testes (Figure 1, H and J). At the end of the first postnatal spermatogenic cycle (34dpp), all germ cells, including fully condensed elongated spermatids, were observed in the seminiferous tubules of *Ip6k1^+/+^*mice but no condensed spermatids were seen in the tubules of 34dpp *Ip6k1^-/-^* mice (Figure 1, K and L). Our histological analysis suggests that *Ip6k1*^-/-^ spermatocytes showed a delay in meiotic progression, but eventually complete meiosis to generate round spermatids. However, as described earlier [5, 10], the development of round spermatids to fully condensed spermatids via the process of spermiogenesis is significantly impaired in the absence of IP6K1.

**Figure 1.**
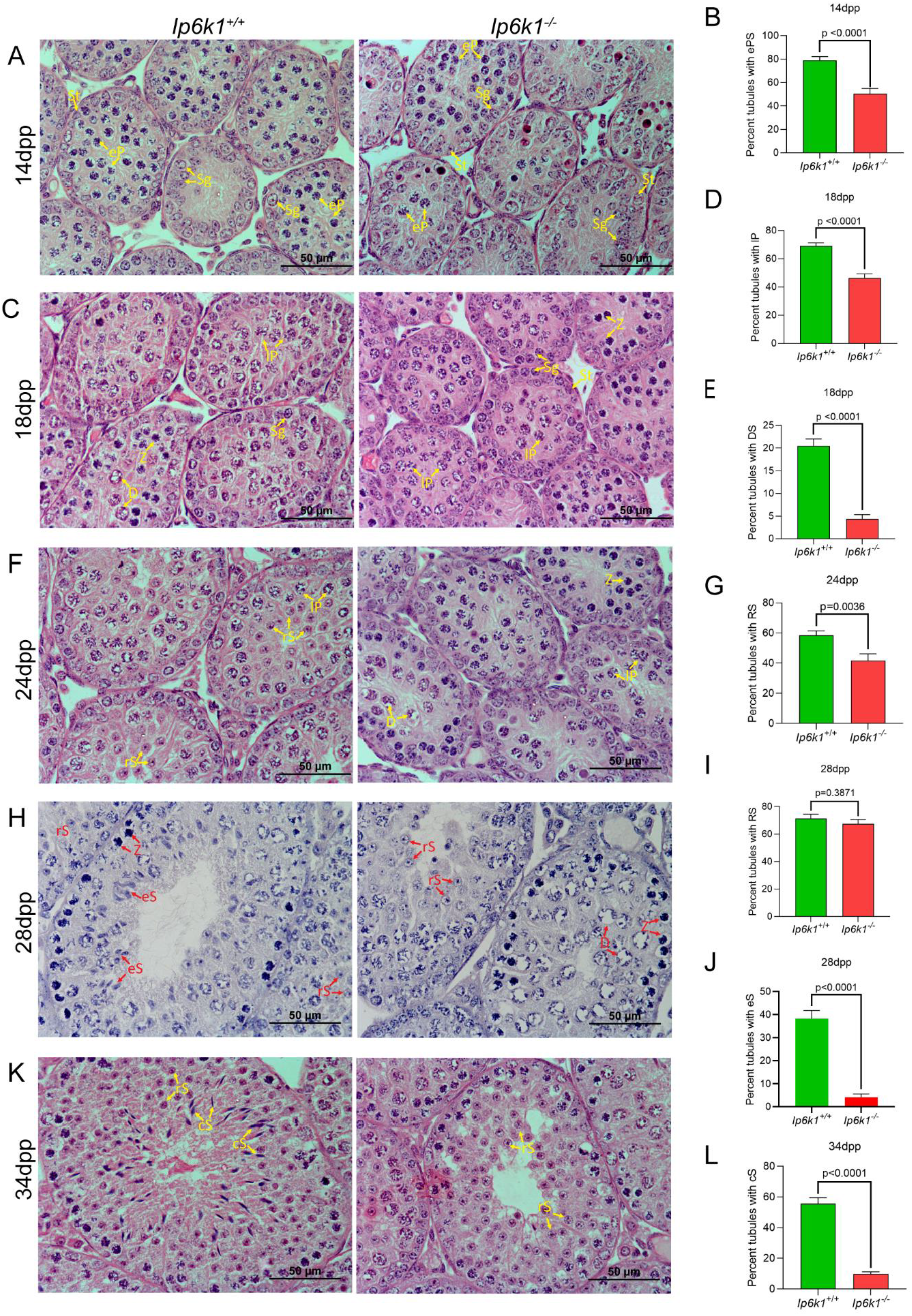
First wave of spermatogenesis is delayed in *Ip6k1* knockout mice. (A) Representative images of H&E stained testes cross-sections of 14dpp *Ip6k1^+/+^* and *Ip6k1^-/-^* mice. (B) Quantification of seminiferous tubules with early spermatocytes in 14dpp testes of two mice of each genotype (n=15 and 11 fields of view for *Ip6k1^+/+^* and *Ip6k1^-/-^* respectively). (C) Representative images of H&E stained testes cross-sections of 18dpp *Ip6k1^+/+^* and *Ip6k1^-/-^* mice. (D and E) Quantification of seminiferous tubules with late pachytene spermatocytes (D) and diplotene spermatocytes (E) in 18dpp testes of three mice of each genotype (n=47 and 48 fields of view for *Ip6k1^+/+^* and *Ip6k1^-/-^* respectively (D), and n=48 and 45 fields of view for *Ip6k1^+/+^* and *Ip6k1^-/-^* respectively (E)). (F) Representative images of H&E stained testes cross-sections of 24dpp *Ip6k1^+/+^*and *Ip6k1^-/-^* mice. (G) Quantification of seminiferous tubules with round spermatids in 24dpp testes of three mice of each genotype (n=24 fields of view for each genotype). (H) Representative images of H&E stained testes cross-sections of 28dpp *Ip6k1^+/+^*and *Ip6k1^-/-^* mice. (I and J) Quantification of seminiferous tubules with round spermatids (I) and elongating spermatids (J) in 28dpp testes of three mice of each genotype (n=30 fields of view for each genotype (I), and n=40 and 42 fields of view for *Ip6k1^+/+^* and *Ip6k1^-/-^* respectively (J)). (K) Representative images of H&E stained testes cross-sections of 34dpp *Ip6k1^+/+^* and *Ip6k1^-/-^* mice. (L) Quantification of seminiferous tubules with condensing spermatids in 34dpp testes of two mice of each genotype (n=46 fields of view for each genotype). Individual cell types are marked in red or yellow letters for clarity. St: sertoli cells; Sg: spermatogonia; Z: zygotene spermatocytes; eP: early pachytene spermatocytes; lP: late pachytene spermatocytes; D: diplotene spermatocytes; rS: round spermatids; eS: elongating spermatids; cS: elongated/condensed spermatids. Scale bars: 50 μm. Data (mean±s.e.m) were analysed using a two-tailed unpaired Student’s *t*-test (P values are indicated above each graph).

### Cell adhesion related genes are downregulated in *Ip6k1^-/-^* testis

To investigate the underlying basis of the delay in the first wave of spermatogenesis in *Ip6k1* knockout mice, we compared the transcriptomes of 17dpp *Ip6k1^+/+^* and *Ip6k1^-/-^*testis by gene expression microarray analysis. Microarray data was analysed by selecting all the genes that showed 1.5-fold or higher upregulation (log base 2 ≥ 0.6) and 1.5-fold or lower downregulation (log base 2 ≤ -0.6), and classified into the following sets: (i) genes downregulated in 17dpp *Ip6k1^-/-^* testis compared with 17dpp *Ip6k1^+/+^*testis (17dpp downregulated), and (ii) genes upregulated in 17dpp *Ip6k1^-/-^*testis compared with 17dpp *Ip6k1^+/+^* testis (17dpp upregulated) (Supplemental Table 1, Figure 2A). The number of downregulated transcripts (∼470) was significantly higher compared with the upregulated transcripts (∼300). Furthermore, the maximum mean downregulation was 12.3 (log base 2), whereas the highest mean upregulation was 4.57 (log base 2). The datasets were analysed by Functional Annotation Clustering for the enriched ‘Gene Ontology’ (GO and ‘Pathways’ terms using the DAVID tool (https://david-d.ncifcrf.gov/) [19]. Significant enrichment scores (ES≥2, P≤0.01) were obtained for the genes that were downregulated in 17dpp *Ip6k1^-/-^*testis (Supplemental Table 1), but not for genes showing upregulated expression. The terms ‘cell-cell junction’ and ‘cell adhesion’ were among the downregulated clusters with significant enrichment scores (Supplemental Table 1, Figure 2B). Interestingly, the same biological processes were seen to be downregulated in our previously reported comparative gene expression analysis of *Ip6k1^+/+^*and *Ip6k1^-/-^* mouse embryonic fibroblasts [9]. We used real time quantitative PCR (RT-qPCR) to confirm the downregulation of some of the transcripts in the 17dpp downregulated set, including some transcripts encoding proteins involved in cell adhesion, such as Cldn3, Cldn7, Cldn8, Gjb3, Cdh1, and Cdh16 (Figure 2, C and D). These genome wide changes in the pathways of cell adhesion are suggestive of a role for IP6K1 in the maintenance of epithelial organisation within the mouse testis.

**Figure 2.**
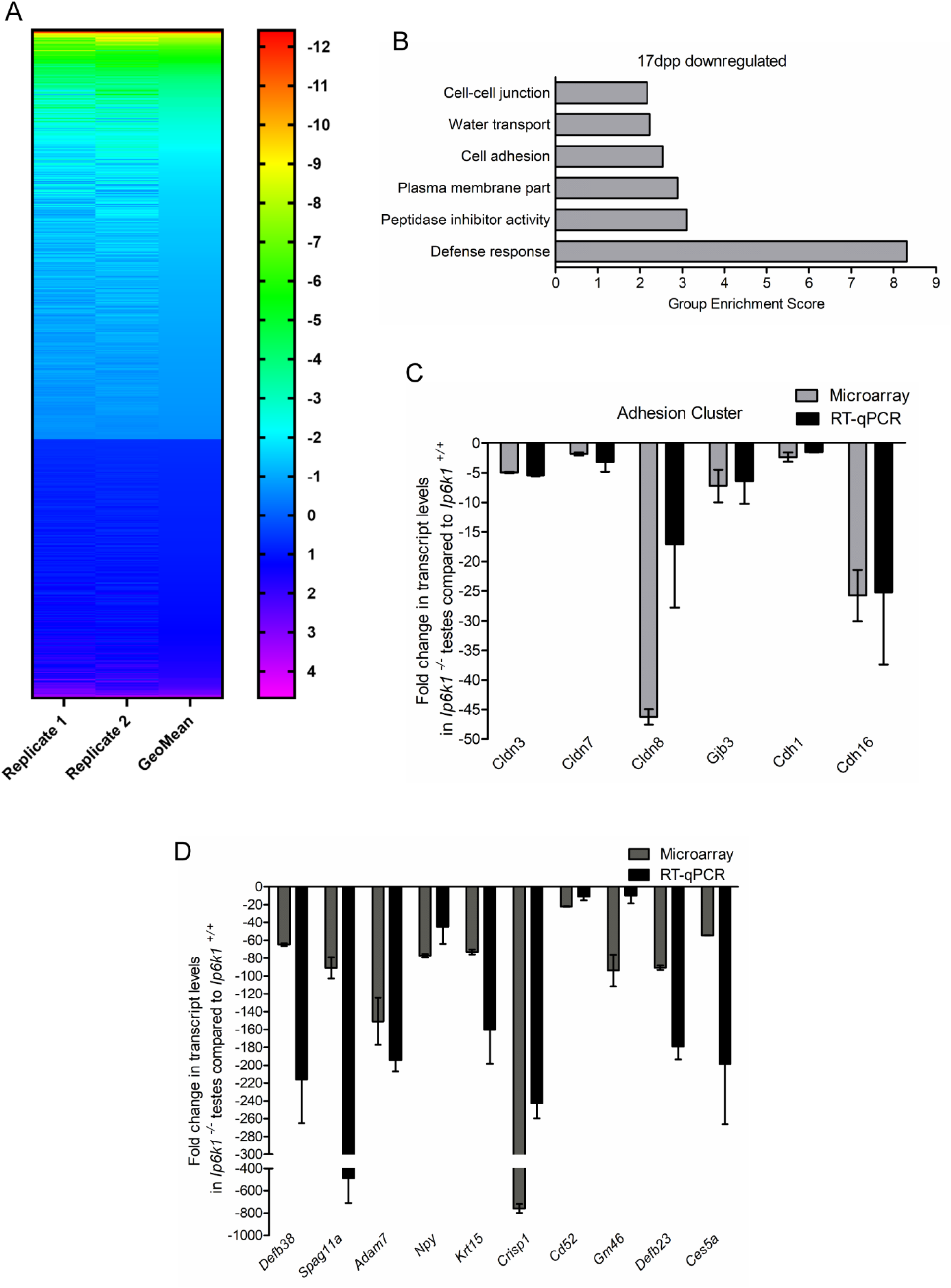
Downregulated expression of cell adhesion genes in *Ip6k1^-/-^* mice testes. (A) Heat map showing change in the expression of gene transcripts that displayed upregulation of log base 2 ≥ 0.6 and downregulation of log base 2 ≤ -0.6 in testis from 17dpp *Ip6k1^-/-^*mice compared to *Ip6k^+/+^* mice. (B) Functional Annotation Clustering was performed for the transcripts downregulated in 17dpp *Ip6k1^-/-^* testes compared with *Ip6k^+/+^*using DAVID tool. Graph depicts enriched terms plotted against group enrichment score (ES≥2). Terms related to cell adhesion were found to be significantly enriched. (C) Validation of microarray data for cell adhesion related transcripts by real-time quantitative PCR. (D) Real-time quantitative PCR analysis of some genes shown to be downregulated by microarray analysis in *Ip6k1^-/-^*testes.

### Loss of IP6K1 causes disruption of the blood-testis barrier in mice

A prior study has demonstrated that the apical ES, which connects elongating spermatids with Sertoli cells, is disrupted in *Ip6k1^-/-^* tubules [11]. This study however did not examine the integrity of the BTB in *Ip6k1^-/-^* testes. Our gene expression analysis revealed that transcripts encoding Claudin family proteins (*Cldn3*, *Cldn7*, and *Cldn8*), which are transmembrane proteins at tight junctions in the BTB, are downregulated in 17dpp *Ip6k1^-/-^* testes (Figure 2C), suggesting that the BTB may be disrupted in the absence of IP6K1. We conducted western blotting analysis to examine the levels of different BTB marker proteins in *Ip6k1^-/-^* testes. In adult mice, we observed an approximately 30% downregulation of claudin 3 expression in knockout testes (Figure 3A). Claudin 3 is known to localize to Sertoli cell-Sertoli cell tight junctions in the BTB, and also to cell membranes of preleptotene/leptotene spermatocytes, aiding the transition of these germ cells across the BTB during meiosis [20, 21]. Immunostaining of testis sections revealed robust claudin 3 staining at the BTB around preleptotene/leptotene spermatocytes in *Ip6k1^+/+^* tubules, whereas weak staining was observed in *Ip6k1^-/-^* tubules (Figure 3B). Conversely, claudin 11, an exclusive Sertoli cell tight junction marker [22] showed similar immunostaining in knockout and wild type tubules, suggesting that all cell adhesion proteins are not dysregulated in *Ip6k1^-/-^*tubules (Figure 3C). We also examined the expression of occludin, another tight junction protein that is known to be essential for maintenance of the BTB [23]. Occludin expression was largely stable in juvenile wild type and knockout testes (Supplemental Figure 2A), but expression of its shorter splice isoform, which appeared later during development, was downregulated in adult knockout testes (Figure 3D).

**Figure 3:**
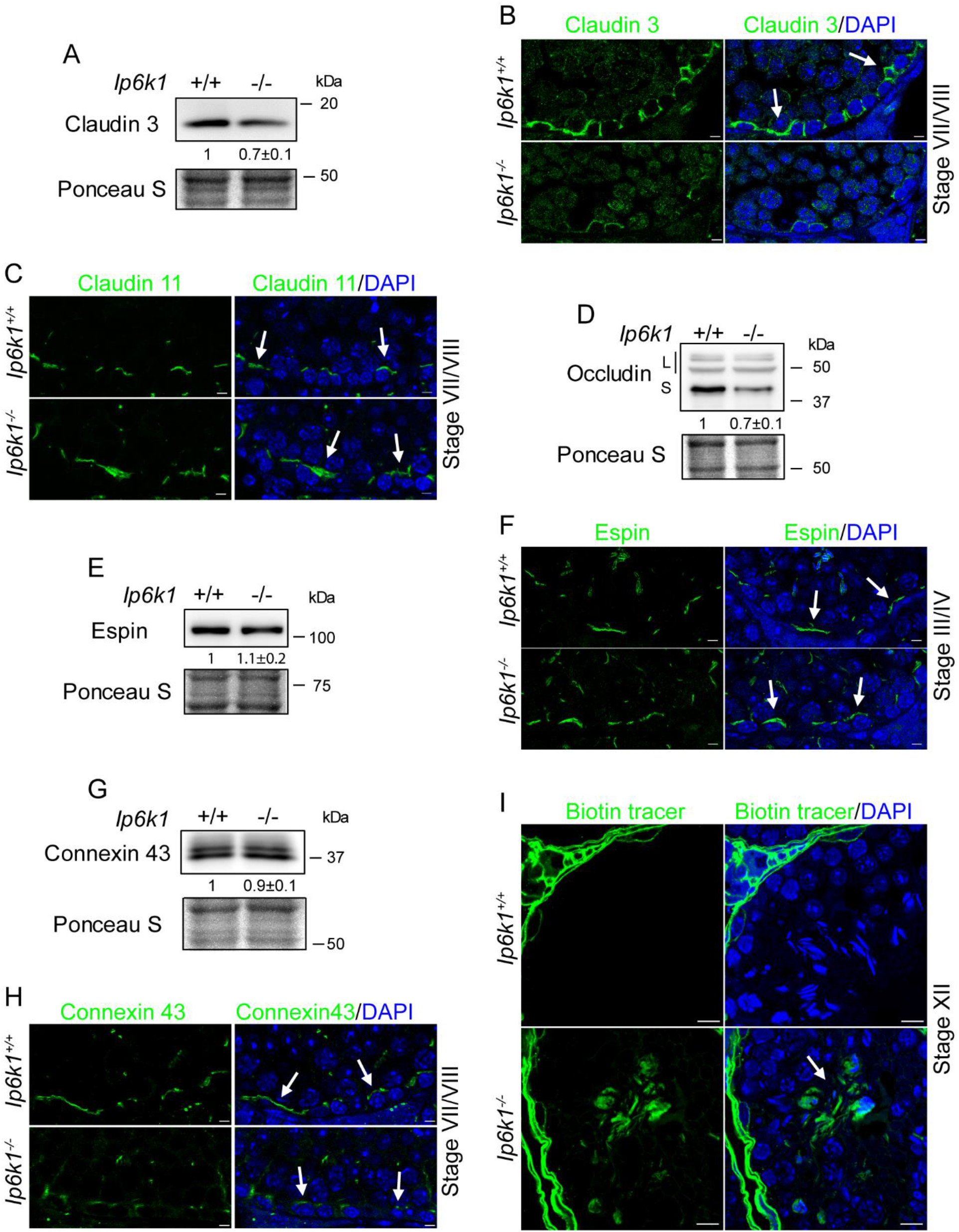
Loss of IP6K1 makes the BTB leaky. (A) Representative immunoblot for tight junction protein claudin 3 in whole testis extracts. Total protein detected by Ponceau S was used as the loading control. The expression of claudin 3 is lower in *Ip6k1^-/-^* testis compared with *Ip6k1^+/+^*. The levels of claudin 3 normalized to the levels of Ponceau S are indicated as the mean±s.e.m. from three biological replicates. (B, C) Immunostaining of adult testis cross-sections of *Ip6k1^+/+^* and *Ip6k1^-/-^*mice to detect claudin 3 and claudin 11 (green). Nuclei are counterstained with DAPI (Blue). Claudin 3 staining is disrupted at *Ip6k1^-/-^* mice BTB compared with *Ip6k1^+/+^* whereas claudin 11 staining seems normal. Scale bars: 5 μm. (D) Representative immunoblot for tight junction protein occludin in whole testis extracts. Total protein detected by Ponceau S was used as a loading control. The expression of the smaller isoform of occludin (S) is lower in *Ip6k1^-/-^*testis compared with wild type. The levels of the larger isoforms of occludin (L) are similar in *Ip6k1^-/-^* testis as compared to wild type. The levels of occludin (S) normalized to the levels of Ponceau S are indicated as the mean±s.e.m. from three biological replicates. (E) Representative immunoblot for espin in whole testis extracts of adult mice. Total protein detected by Ponceau S was used as loading control. The levels of espin are similar in *Ip6k1^-/-^* and *Ip6k1^+/+^* testis. The levels of espin normalized to the levels of Ponceau S are indicated as the mean±s.e.m. from three biological replicates. (F) Immunostaining of adult testis cross-sections of *Ip6k1^+/+^*and *Ip6k1^-/-^* mice to detect espin (green). Nuclei are counterstained with DAPI (blue). Espin staining in the BTB is comparable in *Ip6k1^-/-^*and *Ip6k1^+/+^* testis. Scale bars: 5 μm. (G) Representative immunoblot for gap junction protein connexin 43 in whole testis extracts. Total protein detected by Ponceau S was used as a loading control. The levels of connexin 43 are similar in *Ip6k1^-/-^* and *Ip6k1^+/+^* testes. The protein levels normalized to the levels of Ponceau S are indicated as the mean±s.e.m. from two biological replicates. (H) Immunostaining of adult testis cross-sections of *Ip6k1^+/+^* and *Ip6k1^-/-^* mice to detect connexin 43 (green). Nuclei are counterstained with DAPI. Connexin 43 staining is disrupted at the BTB of *Ip6k1^-/-^* mice compared with *Ip6k1^+/+^*. Scale bars: 5 μm. (I) Staining of adult testis cross-sections of *Ip6k1^+/+^* and *Ip6k1^-/-^* mice to detect EZLink Sulfo-LC-biotin tracer (green). Nuclei are counterstained with DAPI (Blue). The tracer crosses the BTB in *Ip6k1^-/-^* mice testes. Scale bars: 10 μm.

In addition to tight junction markers, we also evaluated the expression and localization of other BTB proteins including espin, an actin bundling protein that marks the ES [24], and connexin 43, which marks gap junctions. Espin expression increased steadily during the development of juvenile wild type testes, reflecting its enrichment in the apical ES (Supplemental Figure 2A). Lower espin levels in 25 and 34 dpp knockout testes (Supplemental Figure 2A) reflect the delay in the appearance of spermatids during the first wave of spermatogenesis (Figure 1). Adult mice however display similar levels of espin in wild type and knockout testes (Figure 3E). This was also reflected in epsin immunostaining, which appeared normal at the basal ES in *Ip6k1^-/-^* tubules (Figure 3F). Expression of the gap junction protein connexin 43 decreased to the same extent during development in *Ip6k1^-/-^*and *Ip6k1^+/+^* juvenile testes (Supplemental Figure 2A), and also showed no significant difference in expression levels between *Ip6k1^-/-^*and *Ip6k1^+/+^* adult testes (Figure 3G). However, as observed for tight junction markers, connexin 43 immunostaining was disrupted at the BTB in *Ip6k1* knockout testes (Figure 3H), suggesting that it may be mislocalized, and revealing that gap junctions at the BTB are also affected by the absence of IP6K1.

The observed alterations in localization and expression of several proteins at the BTB prompted us to examine the integrity of the BTB in *Ip6k1^-/-^* mice. We examined BTB permeability by injecting a biotinylated tracer into the interstitial space between seminiferous tubules in anesthetized mice. Detection with fluorophore-conjugated streptavidin revealed no presence of the biotinylated tracer beyond the basal compartment in wild type testes, whereas significant staining of the tracer was observed in the adluminal compartment in *Ip6k1^-/-^* testes, confirming that the integrity of the BTB is indeed compromised in the absence of IP6K1 (Figure 3I). Together, these results indicate that IP6K1 protects the architecture of the seminiferous epithelium by regulating the expression and localization of key components of the BTB.

### Absence of IP6K1 results in premature sloughing of round spermatids

Disruption of the BTB has been shown to be accompanied with germ cell apoptosis and exfoliation [14, 23, 25]. We therefore examined whether breakdown of the BTB in the absence of IP6K1 affects the integrity of the seminiferous epithelium. Hematoxylin and eosin (H&E) staining of *Ip6k1^-/-^* testes cross-sections showed that germ cells are loosely attached within the tubules and are also sloughed off into the lumen. (Figure 4A, B). Staining of epididymis sections from *Ip6k1^+/+^* and *Ip6k1^-/-^* mice with the round spermatid marker mouse homolog of vasa (MVH), revealed that round spermatids are exfoliated from the testicular epithelium to the epididymis in *Ip6k1^-/-^* mice (Figure 4C). This premature sloughing of postmeiotic germ cells in the tubules of *Ip6k1* knockout mice is in conformity with an earlier report that showed presence of immature and apoptotic postmeiotic germ cells in *Ip6k1^-/-^* epididymis [11]. The histological abnormalities in *Ip6k1^-/-^* testes were more prominent in 6 month old mice. Compared to *Ip6k1^+/+^* controls, 6 month old *Ip6k1^-/-^* testes showed tubular degeneration with marked decline of round spermatids and the appearance of multinucleated giant cells (Figure 4D). These multinucleated degenerating cells in the testes are commonly associated with enhanced apoptosis [26].

**Figure 4:**
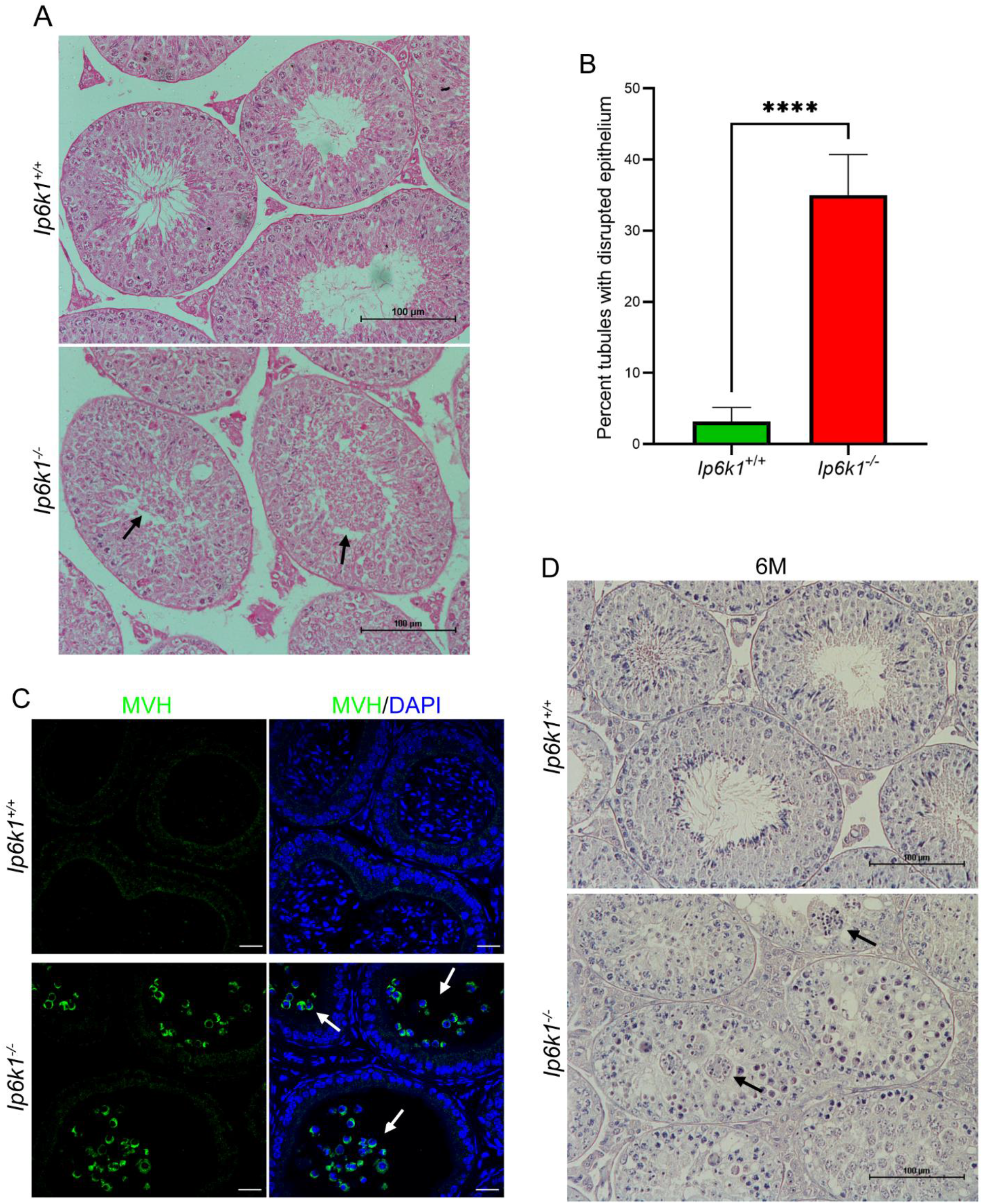
IP6K1 is essential for maintaining the integrity of the seminiferous epithelium. (A) Representative image of H&E stained testes cross-sections of adult *Ip6k1^+/+^*and *Ip6k1^-/-^* mice. Round spermatids are sloughed off into the lumen of the seminiferous tubules. (B) Quantification of seminiferous tubules with disrupted epithelium in three pairs of adult *Ip6k1^+/+^* and *Ip6k1^-/-^* mice. Values indicate mean±s.e.m.; n=13 and 19 for *Ip6k1^+/+^* and *Ip6k1^-/-^* genotypes respectively where n represents the fields of view counted, **** p<0.0001 (two-tailed unpaired Student’s *t*-test). (C) Immunostaining of the round spermatid marker MVH (green) in epididymal cross sections of *Ip6k1^+/+^*and *Ip6k1^-/-^* mice. Arrows indicate MVH-positive round spermatids in *Ip6k1^-/-^* epididymides signifying the premature sloughing off of round spermatids from the testes to epididymides in *Ip6k1^-/-^* mice. Nuclei were counterstained with DAPI (blue). Scale bars: 20 µm. (D) H&E stained testes cross-sections of 6 month old (6M) *Ip6k1^+/+^*and *Ip6k1^-/-^* mice. Multinucleated degenerating giant cells in *Ip6k1^-/-^* mouse testes are indicated by black arrows. Scale bars: 100 μm.

### Absence of IP6K1 promotes disruption of actin-based cytoskeleton in Sertoli cells

The sloughing of round spermatids observed in *Ip6k1^-/-^* testes suggests that there may be defects in the apical ES, in addition to breakdown of the BTB [11]. The apical ES contains transmembrane junctional proteins, including integrins, cadherins and nectins, which connect the apposing plasma membranes of the developing spermatid and the Sertoli cell (Supplemental Figure 1C). These integral membrane proteins are connected with underlying F-actin bundles via various adaptor proteins, including α-actinin. In the brain, IP6K1 has been shown to interact with α-actinin to regulate neuronal migration [27]; IP6K1 may possibly regulate Sertoli cell-germ cell adhesion via a similar mechanism. In the testis, α-actinin forms part of the complex of adaptor proteins that link cadherins with the underlying actin cytoskeleton at the ES [28, 29]. To probe the possibility that IP6K1 acts via α-actinin in the testis, we co-stained testis sections with antibodies directed against both proteins. As reported previously [10], IP6K1 was localized mostly to the cytoplasm of germ cells (Figure 5A), where it did not co-localize with α-actinin (Figure 5B). We observed intense α-actinin staining in the peri-nuclear region and cytoplasm of Sertoli cells in all stages of the seminiferous epithelium, except for stage VII/VIII where adherens junctions are known to turnover as the tubules prepare for spermiation [12] (Figures 5A and Supplemental Figure 2B). To confirm that α-actinin is present in Sertoli cells, we co-stained tubules to detect the intermediate filament protein vimentin, which is known to localize to Sertoli cells [30, 31]. We observed perfect co-localization of α-actinin and vimentin in the cytoplasmic filaments of Sertoli cells (Figure 5, C and D). Surprisingly, despite poor expression of IP6K1 in Sertoli cells, we noted a dramatic reduction in α-actinin staining in Sertoli cells of *Ip6k1* knockout tubules (Figure 5, E and F). At the protein level however, we detected equivalent expression of α-actinin in *Ip6k1^+/+^* and *Ip6k1^-/-^* testes, with expression decreasing over the course of development (Figure 5 G). We therefore conclude that IP6K1 is essential to main the localization of α-actinin in Sertoli cells, but does not appear to influence the levels of α-actinin in the testes.

**Figure 5:**
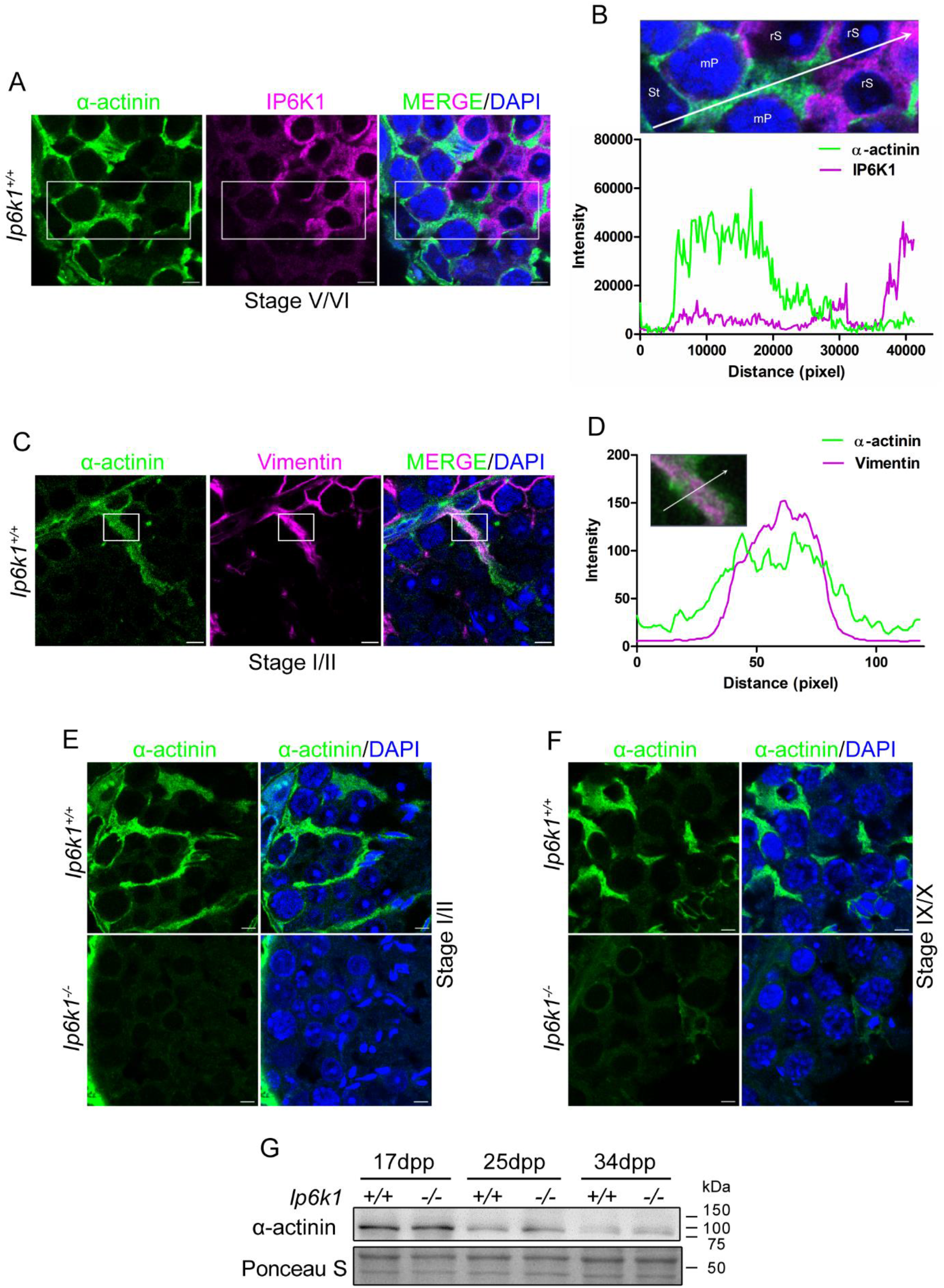
*α*-actinin staining is negligible in the Sertoli cell cytoplasm of *Ip6k1^-/-^* mice. (A) Immunostaining of adult testis cross-sections of *Ip6k1^+/+^* mice to detect α-actinin (green) and IP6K1 (magenta). Nuclei were counterstained with DAPI (blue). Scale bars: 5 μm. (B) Localization analysis of α-actinin (green) and IP6K1 (magenta) in the indicated region (inset) of (A). α-actinin is mainly localized in the cytoplasm of the Sertoli cell whereas IP6K1 is largely detected in the cytoplasm of round spermatids. St: Sertoli cell, mP: mid Pachytene spermatocyte, rS: round spermatids. (C) Immunostaining of adult testis cross-sections of *Ip6k1^+/+^* mice to detect α-actinin (green) and vimentin (magenta). Nuclei were counterstained with DAPI (blue). Scale bars: 5 μm. (D) Localization analysis of α-actinin (green) and vimentin (magenta) in the indicated region (inset) of (C). α-actinin is localized with vimentin in the cytoplasm of the Sertoli cell. (E and F) Immunostaining of adult testis cross-sections of *Ip6k1^+/+^*and *Ip6k1^-/-^* mice at the indicated stages to detect α-actinin (green). Nuclei were counterstained with DAPI (blue). Strong α-actinin staining is seen in the Sertoli cell cytoplasm of *Ip6k1^+/+^*mice but negligible staining is detected in *Ip6k1^-/-^* mice. Scale bars: 5 μm. (G) Representative immunoblot for α-actinin in whole testes extracts of *Ip6k1^+/+^* and *Ip6k1^-/-^* juvenile mice of indicated age (days postpartum; dpp). Total protein detected by Ponceau S was used as a loading control. α-actinin expression decreases with age in juvenile mice.

### Actin cytoskeleton disruption is associated with upregulated integrin and growth factor receptor signalling

Due to the low level of co-expression of IP6K1 and α-actinin, a direct role for IP6K1 in the regulation of germ cell-Sertoli cell adherens junctions via binding to α-actinin is unlikely. We therefore wondered whether IP6K1 indirectly regulates cell signalling pathways that influence cell adhesion. Cell adhesion is regulated by signalling through cell surface junction proteins including integrins and cadherins, and also through growth factor receptor signalling [32-34]. Treatment of rodents with the potential male contraceptive drug adjudin (AF-2364), which induces sloughing of germ cells in seminiferous tubules, has been shown to upregulate expression of β1 integrin, cadherins and catenins, and result in actin filament disruption [35]. We examined the expression of β1 integrin in adult testes, and noted an approximately 40% increase in the level of β1 integrin in *Ip6k1*^-/-^ testes compared with *Ip6k1*^+/+^ testes (Figure 6A). Other adherens junction markers, the cadherin-bound adaptor protein β-catenin, and the immunoglobulin-like adhesion protein nectin-3, were also upregulated in knockout testes (Figure 6, B and C). These data show that, in a manner similar to treatment with adjudin, germ cell sloughing exhibited by *Ip6k1^-/-^* testis is associated with increased expression of adherens junction proteins.

**Figure 6:**
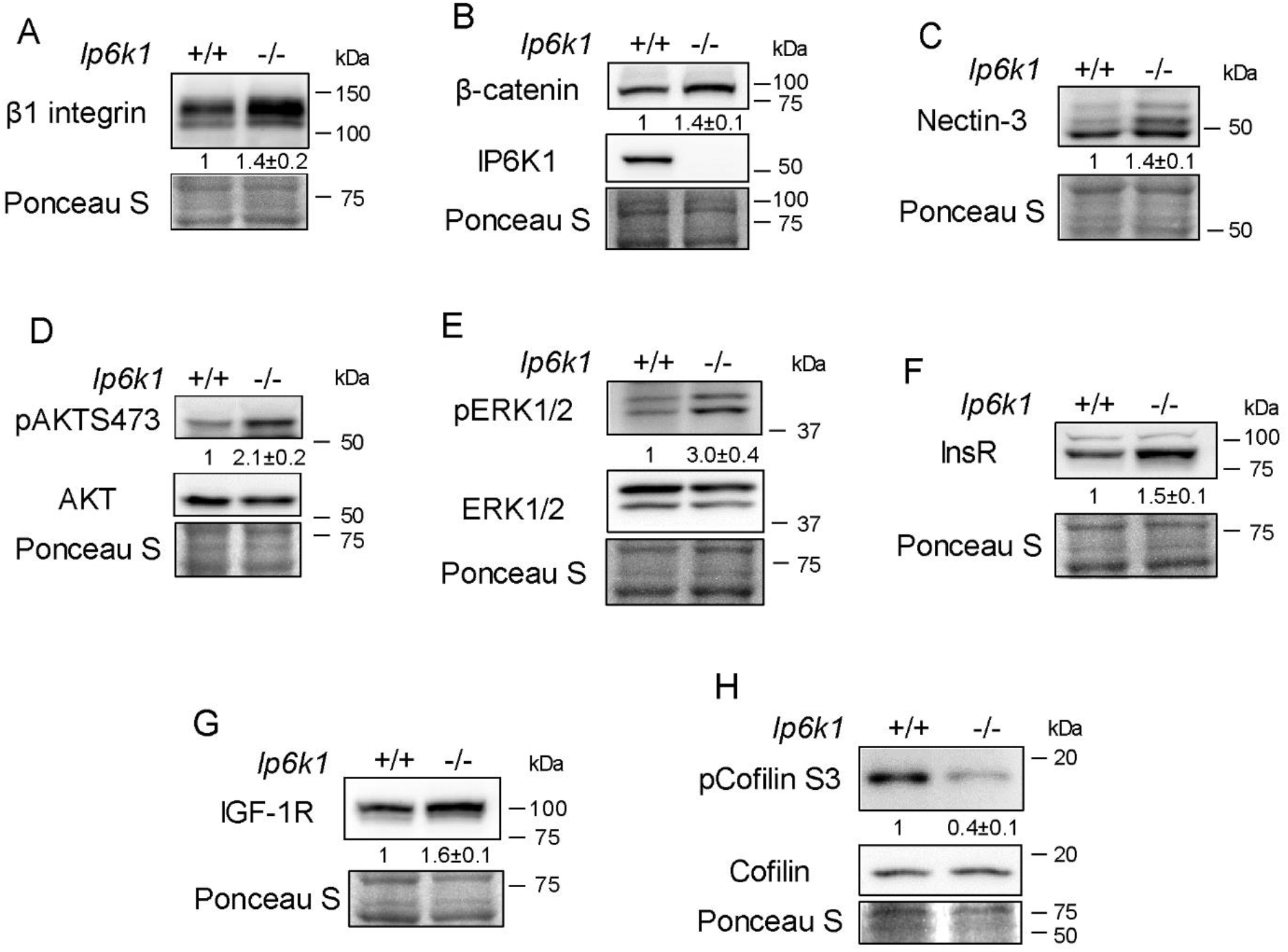
Growth factor receptor and cell junction signalling is deregulated in *Ip6k1^-/-^* testis. (A-C) Representative immunoblots for the detection of β1 integrin (A), β-catenin (B), and nectin-3 (C) and in whole testis extracts. Total protein detected by Ponceau S was used as a loading control. The levels of each protein in *Ip6k1^-/-^* compared with *Ip6k1^+/+^* testes are indicated as the mean±s.e.m. from two (C) or three (A and B) biological replicates. The expression of all three proteins is higher in *Ip6k1^-/-^* testis compared with *Ip6k1^+/+^*. (D and E) Representative immunoblots for the detection of phosphoAKT (pAKTS473) and AKT (D) or phosphoERK1/2 (pERK1/2) and ERK1/2 (E) in whole testis extracts. The levels of the phosphoprotein normalized to the corresponding total protein, in *Ip6k1^-/-^* testis compared with *Ip6k1^+/+^*, are indicated as the mean±s.e.m. from two (E) or three (D) biological replicates The phosphorylation of AKT and ERK1/2 is higher in *Ip6k1^-/-^* testis compared with *Ip6k1^+/+^*. (F and G) Representative immunoblots for the detection of insulin receptor (InsR; F) and insulin like growth factor 1 receptor (IGF-1R; G) in whole testis extracts. Total protein detected by Ponceau S was used as a loading control. The expression of InsR is higher in *Ip6k1^-/-^* testis compared with *Ip6k1^+/+^*. The levels of each protein in *Ip6k1^-/-^*compared with *Ip6k1^+/+^* testes are indicated as the mean±s.e.m. from three biological replicates. The expression of both proteins is higher in *Ip6k1^-/-^* testis compared with *Ip6k1^+/+^*. (H) Representative immunoblot for the detection of phospho-cofilin (pCofilin) and cofilin in whole testis extracts. The phosphorylation of cofilin is lower in *Ip6k1^-/-^* testis compared with *Ip6k1^+/+^*. The levels of the phosphoprotein normalized to the corresponding total protein, in *Ip6k1^-/-^* testis compared with *Ip6k1^+/+^*, are indicated as the mean±s.e.m. from three biological replicates.

Adjudin treatment also leads to increased signalling via the β1 integrin/PI3K/AKT/ERK pathway in Sertoli cells [35, 36]. 5-IP7 synthesized by IP6K1 has been shown to downregulate AKT signalling by competing with PI(3,4,5)P3 to bind the PH domain of AKT and preventing AKT localization at the plasma membrane [37]. Despite the low level of IP6K1 expressed in Sertoli cells (Figure 5B), it is possible that a decrease in 5-IP7 in the absence of IP6K1 promotes activation of AKT. We therefore examined whether signalling via the AKT pathway is upregulated in *Ip6k1* knockout testes by monitoring the level of phosphorylated AKT. Phosphorylation of AKT on Ser 473 was two-fold higher in *Ip6k1*^-/-^ testes compared with wild type (Figure 6D). Additionally, phosphorylation of ERK1/2 was also significantly upregulated in the absence of IP6K1 (Figure 6E).

Another pathway upstream to AKT and ERK activation, which is known to regulate cell adhesion, is growth factor receptor signalling via insulin / insulin-like growth factor (IGF) receptors [33]. Studies in somatic cells have shown that insulin treatment promotes disruption of actin filaments, especially under conditions of high insulin receptor expression [38-40]. We noted a 50-60% increase in the expression of insulin receptor and IGF-1 receptor in *Ip6k1*^-/-^ testes compared with *Ip6k1*^+/+^ testes (Figure 6, F and G). This upregulation of insulin signalling may also contribute to increased AKT activation in the absence of IP6K1. It has been shown that increased signalling through the insulin/AKT/MEK axis leads to activation of cofilin, an actin binding protein that stimulates the severance and depolymerisation of actin [41-44]. Cofilin is inactivated by phosphorylation on Ser3, and is activated when this residue is dephosphorylated-this phosphorylation/dephosphorylation of cofilin is a convergence point for adhesion and growth factor signalling pathways that regulate actin cytoskeletal dynamics [44]. We therefore examined whether a decrease in cofilin phosphorylation accompanies the disruption of actin-based filaments observed in *Ip6k1^-/-^* testes. Indeed, we observed a substantial reduction in the extent of cofilin phosphorylation, with no change in cofilin levels in *Ip6k1^-/-^* testes compared with wild type (Figure 6H). In summary, our data suggests that disruption of the Sertoli cell actin cytoskeleton, marked by the loss of α-actinin in *Ip6k1*^-/-^ tubules, is the result of a conjunction of signalling pathways, including cell junction and growth factor receptor signalling, impinging on AKT, ERK and cofilin activation.

## Discussion

Earlier studies have revealed that IP6K1 expression in the testis is essential for condensation of elongating spermatids [10], and to maintain the apical ES [11]. We now reveal another important role for IP6K1 in male fertility - maintaining the functional integrity of the BTB. We observed that loss of IP6K1 leads to downregulated expression and mis-localization of junction proteins found at the BTB, resulting in increased BTB permeability. In contrast, we noted upregulated expression of adherens junction proteins and growth factor receptors, correlating with increased activation of the AKT/ERK signalling pathway. The subsequent downstream activation of cofilin, and disruption in the Sertoli cell actin cytoskeleton, results in loss of germ cell–Sertoli cell adhesion, ultimately leading to premature exfoliation of round spermatids in *Ip6k1*^-/-^ mice (Figure 7).

**Figure 7.**
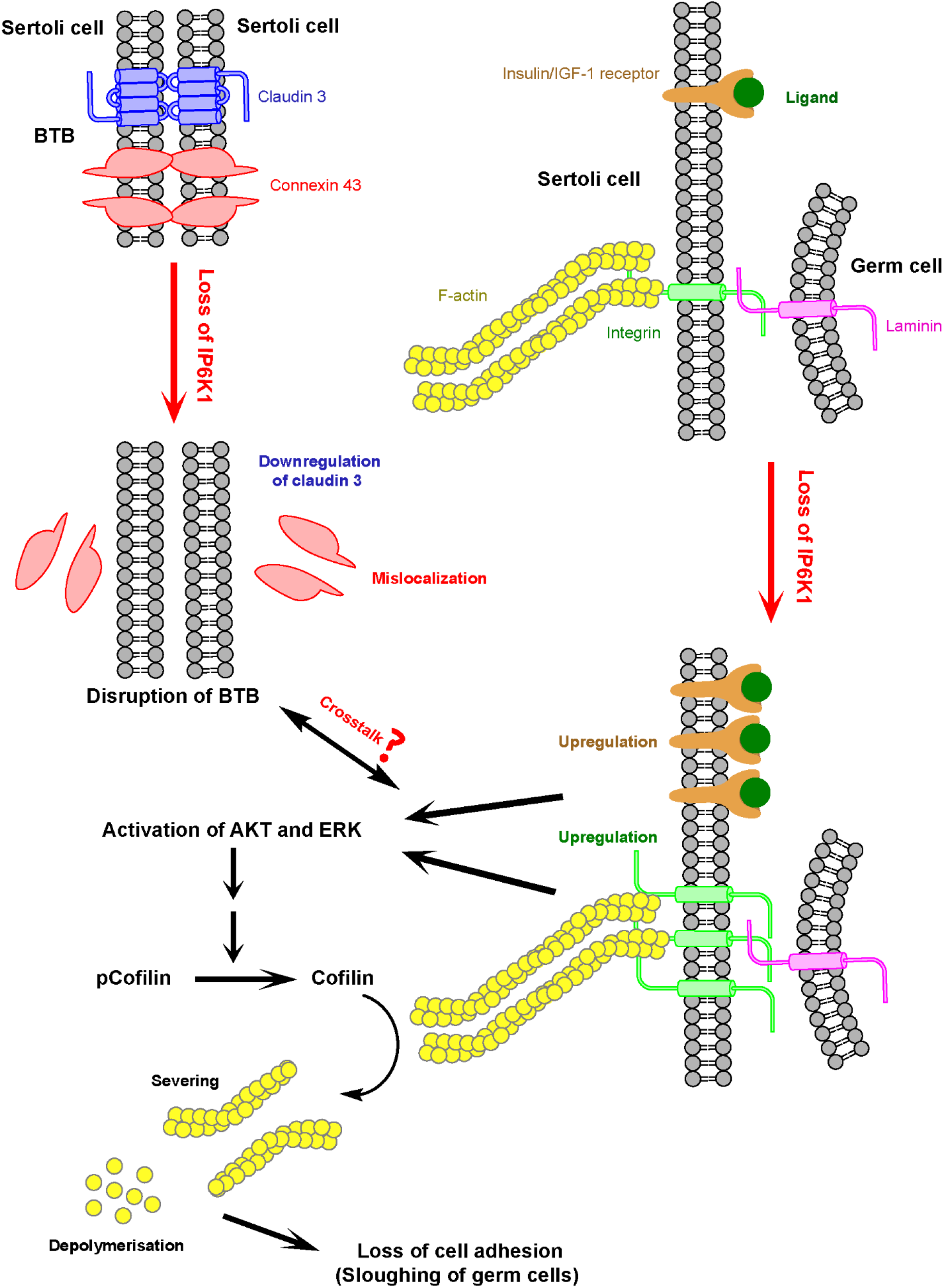
Model depicting the underlying basis of disruption of cell junctions in *Ip6k1^-/-^* mouse testis. Loss of IP6K1 promotes disruption of both the BTB and the actin based cytoskeleton in Sertoli cells. Disruption of the BTB involves both mislocalization and downregulation of its constituent proteins. On the other hand, disruption of actin based cytoskeleton in the Sertoli cells is associated with the upregulation of insulin/IGF-1 receptors and β1 integrin, and downstream activation of AKT, ERK and cofilin.

Gene expression analysis in testes from 17 day old mice revealed large-scale transcription dysregulation in the absence of IP6K1. Our earlier work has shown that mouse embryonic fibroblasts lacking IP6K1 also show global changes in gene expression [9]. Interestingly, in both studies, we noted a marked downregulation of genes involved in cell adhesion pathways. There are several possible ways by which IP6K1 could regulate gene expression. We have earlier shown that IP6K1 interacts with protein complexes on the mRNA cap to promote translational suppression and upregulate the formation of processing bodies [45]. This could in turn have a cascading effect on gene expression. IP6K1 may also interact with proteins that directly regulate mRNA transcription, export, or stability. 5-IP7 synthesized by IP6K1 may regulate proteins involved in mRNA metabolism, either via serine pyrophosphorylation or direct binding to proteins. IPMK, an inositol phosphate kinase that is structurally related to IP6Ks, has been shown to act directly as a transcriptional co-activator [46]. Transcription may also be regulated via chromatin remodelling – inositol pyrophosphates have been shown to regulate histone deacetylase and histone demethylase activities in yeast and mammals [47, 48]. As IP6 is known to regulate the export of nascent transcripts from the nucleus to the cytoplasm [49, 50], the loss of IP6K1 could alter IP6-dependent mRNA export.

Our study shows that IP6K1 is required for the assembly of tight junctions and gap junctions at the BTB, and is also essential for proper assembly of actin filaments at adherens junctions between germ cells and Sertoli cells. It is not clear whether the loss of IP6K1 promotes the disassembly of various types of junctions independent of each other, or whether there is cross-talk between disruption of the BTB and germ cell–Sertoli cell junctions in the testis of *Ip6k1*^-/-^ mice. Disassembly of one type of junction in the seminiferous epithelium has been shown to affect the stability of other types of junctions [25, 51, 52]. Disruption of gap junctions upon injection of a connexin mimic peptide is known to promote mislocalization of the adherens junction marker N-cadherin, and decrease the levels of the tight junction protein occludin [25]. Similarly, in *Ip6k1*^-/-^ tubules we observe concomitant disruption of the tight junction marker claudin 3, and the gap junction marker connexin 43 at the BTB, along with loss of the actin cytoskeleton marker α-actinin in Sertoli cells.

All three IP6 kinases have been implicated in the regulation of cell adhesion and actin cytoskeleton architecture. The loss of IP6K2 in HCT116 colorectal carcinoma cells causes defects in cell adhesion and migration, leading to reduced ability of these cells to form tumours in immunocompromised mice [53]. This effect is due to nuclear sequestration and inactivation of liver kinase B1 by 5-IP7 synthesized by IP6K2 [53]. Deletion of IP6K3 in mice alters cytoskeletal architecture in cerebellar Purkinje cells, leading to defects in motor learning and coordination [54]. IP6K3, but not the other two IP6Ks, was shown to directly interact with the actin binding proteins spectrin and adducin, promoting their mutual interaction. In migrating SH-SY5Y neuroblastoma cells, IP6K3 is enriched at the leading edge, where it associates with the motor protein dynein to promote phosphorylation of focal adhesion kinase (FAK) and turnover of focal adhesions [55]. This molecular function of IP6K3 has been implicated in the loss of neuronal cell migration and brain malformations in *Ip6k3* knockout mice [55]. IP6K1 has been shown to regulate adhesion and migration in a variety of cell types. Mouse embryonic fibroblasts lacking IP6K1 showed reduced cell spreading and migration, along with decreased phosphorylation of FAK and paxillin [9]. Cancer cell lines deficient in IP6K1 displayed reduced anchorage independent growth, migration and invasion [9]. The importance of IP6K1 in promoting cell invasion was apparent in the lower incidence of oral carcinogen-induced invasive carcinoma in *Ip6k1* knockout mice compared with their wild type counterparts. Mice lacking IP6K1 also show brain malformations and defects in neuronal migration [27]. This was attributed to the binding of IP6K1 with α-actinin, which in turn binds to FAK at the focal adhesion complex. 5-IP7 synthesized by IP6K1 enhances FAK autophosphorylation to promote neuronal migration. Snyder and co-workers have suggested that the interaction between IP6K1 and α-actinin may be responsible for maintaining the apical ES in the mouse seminiferous epithelium [11]. However, our data suggests minimal co-localization of IP6K1 and α-actinin in seminiferous tubules (Figure 5B). Instead, we suggest that increased signalling via the AKT/ERK pathway, mediated by insulin/IGF-1 receptor upregulation, may be the molecular basis for disruption of the ES observed in *Ip6k1* knockout mice. We conclude that a complex interplay of different pathways regulated by IP6K1 is responsible for maintenance of the BTB, germ cell–Sertoli cell adhesion, and development of spermatids in the mouse testis.

## Materials and Methods

### Mice

C57BL/6 *Mus musculus* strain was used for our studies. The mice were housed in the Experimental Animal facility, Centre for DNA Fingerprinting and Diagnostics, Hyderabad, India. All animal experiments were performed in compliance with guidelines provided by the Committee for Control and Supervision of Experiments on Animals, Government of India. All animal experiments were approved by the Institutional Animal Ethics Committee (Protocol number PCD/CDFD/02 – version 2). Generation of the *Ip6k1* gene knockout mouse has been described previously [5]. *Ip6k1* wild type and knockout littermates were produced by breeding *Ip6k1* heterozygous mice. Testes for experimentation were obtained from juvenile, adult (2-3 months old) and 6 months old male mice.

### Reagents and antibodies

All chemicals were procured from Merck, unless specified otherwise. Primary antibodies used for immunoblotting (IB) and immunofluorescence (IF) along with their dilutions were: anti-IP6K1 (GeneTex, GTX103949; 1:3000 IB, 1µg IP); anti-IP6K1 (Sigma, HPA040825; 1:400 IF); anti-β1 integrin (Santa Cruz Biotechnology, sc-6622; 1:1000 IB); anti-α-actinin (Santa Cruz Biotechnology, sc-17829; 1:50 IF, 1:1000 IB); anti-vimentin (Abcam, ab92547; 1:200 IF); anti-occludin (Abcam, ab31721; 1:2500 IB); anti-claudin 11 (Abcam, ab53041; 1:400 IF); anti-claudin 3 (Abcam, ab15102; 1:100 IF, 1:1000 IB); anti-connexin 43 (Abcam, ab11370; 1:1000 IF, 1:5000 IB); anti-nectin-3 (Abcam, ab16913; 1:2500 IB); anti-espin (BD Biosciences, 611656; 1:400 IF, 1:2500 IB); anti-phospho-ERK1/2 (CST, 9106S; 1:700 IB); anti-ERK1/2 (CST, 9102; 1:1000 IB); anti-β-catenin (Invitrogen, 13-8400; 1:2500 IB); anti-MVH (Abcam, ab13840; 1:300 IF); anti-insulin receptor β (CST, 8338T; 1:2500 IB); anti-IGF-I receptor β (CST, 8338T; 1:2500 IB); anti-phospho-AKT S473 (CST, 4060P; 1:1500 IB); anti-AKT (CST, 9272S; 1:1500 IB); anti-phospho-cofilin-S3 (St Johns Laboratory, STJ90230; 1:2000 IB); anti-cofilin (CST, 5175T; 1:2000 IB).

### Staging of the seminiferous epithelium

The staging of the seminiferous epithelium was done based on the presence, morphology and location of various cells types within the seminiferous epithelium. Since the elongated spermatids in *Ip6k1^-/-^*testes do not progress beyond step 11 [10], the condensation pattern of these spermatids was not considered for staging beyond this step. Stages I to VI were identified based on the types of spermatogonia. Stage VII/VIII tubules in *Ip6k1^+/+^* testes were determined by the presence of a lining of elongated spermatids along the lumen of the seminiferous tubule. Since the elongated spermatids degenerate at this stage in *Ip6k1^-/-^* testes, stage VII/VIII tubules in the knockout mice are marked by a negligible number of degenerating spermatids and the presence of densely stained preleptotene spermatids at the base of the seminiferous epithelium. Stage IX/X tubules were determined by the absence of round spermatids and the presence of early elongating spermatids. Stage XI tubules were distinguished from stage IX/X tubules by the presence of thin nuclei with dense chromatin. Stage XII tubules were identified by the presence of meiotic bodies and secondary spermatocytes. The methodology followed for the staging has been described previously [56, 57].

### Histology

Juvenile, adult (2-3months old) and 6 months old mice were used for histological analysis of testes, which was performed as described previously [10]. Briefly, mice were euthanized, testes were dissected, fixed in Bouin’s solution and embedded in paraffin. 4 µm sections were prepared on glass slides, deparaffinised in xylene, rehydrated in a graded series of ethanol, and stained with H&E. Images were obtained using a bright-field light microscope (Nikon ECLIPSE Ni-U, NIS Elements acquisition software, 20X 0.5 N.A. or 40X 0.75 N.A. objectives).

### Tissue immunofluorescence

Paraffin-embedded testis sections were deparaffinised by heating at 60°C for 1-2 h and placing in xylene. Rehydration was done using an ethanol gradient series with a gradual increase in water content. For antigen retrieval, the sections were heated in sodium citrate buffer (10 mM, pH 6) in a microwave oven for 20 min and kept in the solution until it cooled to room temperature. The slides were washed in PBS, permeabilised in 0.5% Triton X-100 for 10-15 min, blocked in blocking buffer (5% BSA+0.1% Triton X-100) for 1 h at room temperature and incubated overnight at 4°C with primary antibodies diluted in blocking buffer. The slides were washed again with PBS and stained with fluorophore-conjugated secondary antibodies for 1 h at room temperature. Alexa Fluor 488-conjugated goat anti-rabbit-IgG, Alexa Fluor 568-conjugated donkey anti-rabbit-IgG, Alexa Fluor 488-conjugated goat anti-mouse-IgG, and Alexa Fluor 488 conjugated donkey anti-goat-IgG secondary antibodies were from Life Technologies. All secondary antibodies were used at 1:400 dilution in blocking buffer. The slides were mounted in antifade mounting medium containing DAPI (H1200, Vector Labs) and sealed with nail polish. Images were acquired with an LSM 700 (Zen acquisition software) confocal microscope (Zeiss) or with Leica SP8 (Leica acquisition software) equipped with 405, 488 and 561 nm lasers, and fitted with a 63X 1.4 NA objective.

### Gene expression microarray

Testes were collected in biological replicates from 17dpp *Ip6k1^+/+^*and *Ip6k1^-/-^* mice, two animals of each age and genotype, under RNase free conditions and immediately frozen in liquid nitrogen. Total RNA was extracted from frozen tissues using the RNeasy Mini Kit (Qiagen, Cat #74106). Further sample processing and microarray analysis was conducted at Genotypic Technologies Pvt. Ltd. Bangalore. The samples were labelled using Agilent Quick-Amp labelling Kit (Cat. # 5190-0442). 500 ng of each sample was incubated with reverse transcription mix at 40°C and converted to double stranded cDNA primed by oligonucleotide dT with a T7 polymerase promoter. cRNA was generated by in vitro transcription of the double stranded cDNA at 40°C, and the dye Cy3 CTP (Agilent) was incorporated during this step. 600 ng of Cy3 labelled RNA samples were fragmented and hybridized using Agilent’s Gene Expression In situ Hybridzation Kit (Cat. # 5190-6420). Hybridization onto Agilent-014868 Whole Mouse Genome Microarray 8×60K (G4858A; AMADID, 28005) was carried out in Agilent’s Surehyb Chambers at 65°C for 16 h. The hybridized slides were washed using Gene Expression wash buffers (Agilent), and scanned using the Agilent Microarray Scanner (G2505C) at 5 μm resolution. Data was extracted using Feature Extraction software and normalized using GeneSpring GX 12.6 Software (75th percentile shift method). Data are presented as the average fold change in gene expression in the two *Ip6k1^-/-^* mice compared with the average expression for that gene in two *Ip6k1^+/+^* mice for each age group and genotype. Gene expression data are available at GEO (GSE153500). The lists of up-and downregulated genes were subjected to Functional Annotation Clustering using the Database for Annotation, Visualization and Integrated Discovery (DAVID) v6.7 tool [19]. Transcripts that showed 1.5-fold up or down regulation were considered for Gene Ontology (GO) term enrichment.

### Real-time quantitative PCR

Validation of microarray data was done by reverse transcription real-time quantitative PCR (RT-qPCR) as previously described [10]. Briefly, total cellular RNA was isolated from 17dpp *Ip6k1^+/+^* and *Ip6k1^-/-^*testes using TRIzol reagent (Invitrogen) and RNeasy Mini Kit (Qiagen) according to the manufacturers’ guidelines. cDNA was synthesised with SuperScript Reverse Transcriptase III (Invitrogen) using oligonucleotide dT primers. qPCR was carried out in ABI 7500 real-time PCR system (Applied Biosystems) using MESA GREEN qPCR MasterMix Plus for SYBR® Assay Low ROX (Eurogentec) for detection. The gene-specific primers used in qPCR reactions are listed in Supplemental Table 2.

### Western blotting

Testes from *Ip6k1^+/+^* and *Ip6k1^-/-^* mice were dissected, placed on ice and briefly washed in cold 1X PBS. Tunica albuginea were removed and seminiferous tubules were minced in RIPA buffer (50 mM Tris-HCl pH 7.4, 150 mM NaCl, 1% Triton X-100, 0.5% sodium deoxycholate, 0.1% SDS, 1 mM EDTA) containing freshly added protease inhibitors (P8340, Sigma-Aldrich), phosphatase inhibitor cocktail 3 (P0044, Sigma-Aldrich), and tyrosine phosphatase inhibitor (sodium orthovanadate, 1 mM; Sigma, S6508). The tissues were subjected to homogenisation (D-1 Homogenizer-Disperser, Miccra) and placed on end over mixing for 1h. The extracts were centrifuged at 15,000 g for 10 min at 4°C and supernatants were separated. A few microliters of the supernatant were kept aside for quantification by BCA assay and the rest was mixed with 4X SDS sample buffer to a final dilution of 1X and heated at 95°C for 5 min. The samples were resolved on SDS-PAGE and transferred onto activated PVDF membrane (GE Healthcare). The PVDF membrane was blocked with 5% non-fat milk (Rockland) for 1-2h at RT, followed by incubation with the protein specific primary antibody overnight at 4°C. The membrane was washed three times with TBST buffer (20 mM Tris-HCl pH 7.6, 150 mM NaCl and 0.1% Tween 20), and incubated at RT for 1 h with the following HRP-conjugated secondary antibodies: goat anti-rabbit IgG conjugated to HRP (Southern Biotech, 4010-05; 1:5000), and goat anti-mouse IgG conjugated to HRP (Southern Biotech, 1031-05; 1:10,000). The membrane was again washed three times with TBST and the protein of interest was detected using a chemiluminescence reagent (ECL Prime, GE Healthcare). Chemiluminescence was detected using the ImageQuant LAS500 imager (GE Healthcare). If required blots were stripped using Restore™ Western Blot Stripping Buffer (ThermoFisher Scientific), and re-probed with another antibody. Densitometry analysis of bands was performed by using Image Studio Lite Ver 5.2 (LI-COR Biosciences). The band intensity of the protein of interest was normalised to total protein detected by Ponceau S.

### BTB Permeability assay

Mice were anaesthetized by injecting Ketamine (20-40 mg/kg body weight) and Xylazine (2mg/kg body weight) intraperitoneally. Under aseptic conditions a 1–1.5 cm incision was made anterior to the penis, to cut through skin and muscle. Testes were removed from the scrotal sac with curved sterile forceps through the incision by gently pulling the dorsal fat pad associated with the testes. 50 µl of freshly prepared solution of Biotin tracer (10 mg/ml EZ-Link Sulfo-NHS-LC-Biotin, Thermo Fisher Scientific, in PBS containing 1 mM CaCl2) was injected slowly into the inter-tubular space of one testis using a 30-gauge needle, and the other testis was injected similarly with PBS-CaCl2 solution which lacked the Biotin tracer. Mice were placed in a recovery chamber; after 30 min, while the mice were still under anaesthesia, they were euthanized. The testes were dissected, fixed, and processed for histological examination to detect biotin using fluorescently tagged streptavidin (Streptavidin, Alexa Fluor^TM^ 488 Conjugate, S32354).

### Statistical Analyses

Images shown in the figures are representative. The number of animals used (n) to obtain the quantitative data is mentioned in the figure legends. Data was analysed using Image Studio Lite Ver 5.2 (LI-COR Biosciences) and GraphPad Prism software.

## Acknowledgements

We acknowledge the staff of the CDFD experimental animal facility and microscopy core facility. We thank all members of the Laboratory of Cellular Signalling for helpful comments.

## Author contributions

SAB: Conceptualization, Methodology, Investigation, Validation, Formal Analysis, Visualization, Writing-Original Draft and Editing.

ABM: Conceptualization, Methodology, Investigation, Validation, Visualization, Writing-Review and Editing.

VO: Investigation (Gene expression analysis, Mouse colony management) JS: Investigation (Assistance with testis histology)

RB: Conceptualization, Supervision, Formal Analysis, Validation, Writing-Review & Editing, Project Administration, Funding Acquisition

