## Supplemental Figures S1-S2 for "Inositol hexakisphosphate kinase 1 is essential for cell junction integrity in the mouse seminiferous epithelium"

**ORCID iDs**

This PDF file includes:

Supplemental Figures S1 and S2

### Supplemental Figure 1

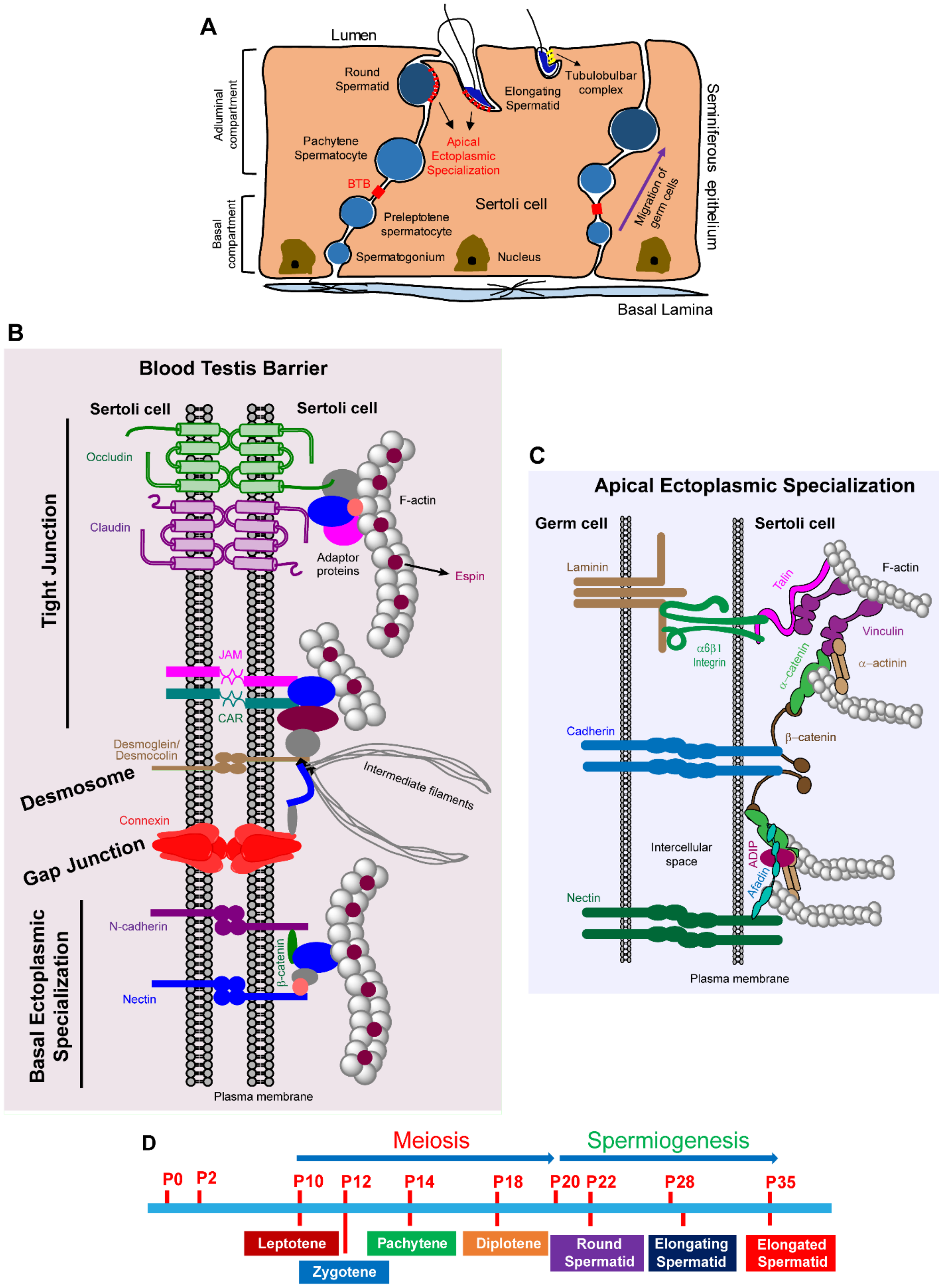

**Supplemental Figure 1.** (A) Diagram showing architecture of the seminiferous epithelium. BTB, which is present between adjacent Sertoli cells, divides the seminiferous epithelium into basal and adluminal compartments. Apical ectoplasmic specialization connects spermatids with the Sertoli cell. (B) Diagram showing the composition of the BTB. It consists of four types of junctions: tight junctions, desmosomes, gap junctions and basal ectoplasmic specialization. Each type of junction is composed of membrane proteins connected to the underlying cytoskeleton via adaptor proteins. (C) Diagram depicting the components of the apical ES. It is an actin based cell junction composed of membrane proteins which are connected to actin filaments via various adaptor proteins. The membrane proteins include laminin that binds integrins, cadherin, and nectin. (D) Diagram depicting various germ cell types that appear at different days postpartum (dpp) in mice testes during the first wave of spermatogenesis.

### Supplemental Figure 2

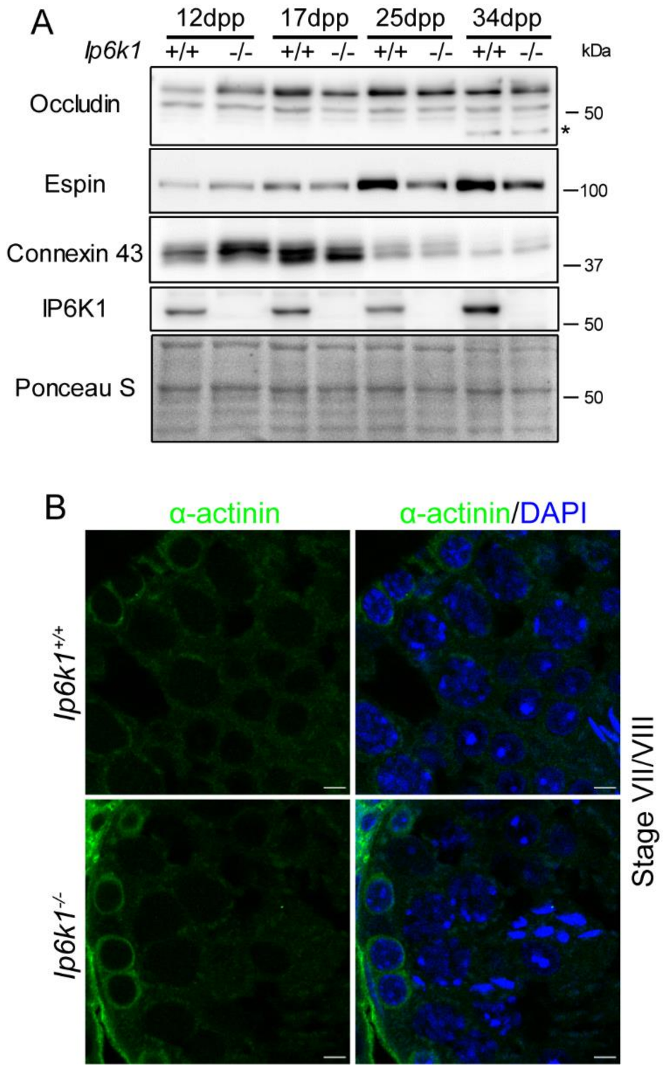


**Supplemental Figure 2.** (A) Representative immunoblots from two experimental sets to detect expression of occludin, espin, and connexin 43 in whole testis extracts of *Ip6k1*$^{+/+}$ and *Ip6k1*$^{-/-}$ juvenile mice of 12, 17, 25, and 34 days postpartum (dpp). Total protein detected by Ponceau S was used as a loading control. The smaller isoform of occludin is marked by an asterisk (*). (B) Immunostaining of adult testis cross-sections of *Ip6k1*$^{+/+}$ and *Ip6k1*$^{-/-}$ mice to detect α-actinin (green) at stage VII/VIII. Nuclei were counterstained with DAPI (blue). α-actinin staining is absent in the Sertoli cell cytoplasm of both *Ip6k1*$^{+/+}$ and *Ip6k1*$^{-/-}$ mice at this stage. Scale bars: 5 µm.
